# PATIENT-DERIVED ORGANOIDS CAPTURE HISTOLOGICAL, MOLECULAR AND THERAPEUTIC HETEROGENEITY IN PHARYNGEAL AND LARYNGEAL SQUAMOUS CELL CARCINOMAS

**DOI:** 10.64898/2026.03.24.713954

**Authors:** Miguel Álvarez-González, Esperanza Pozo-Agundo, Beatriz De Luxán-Delgado, Helena Codina-Martínez, Borja Gallego, María Otero-Rosales, Israel Rivera-García, Andrea Blázquez, Mar Rodríguez-Santamaría, Daniela Corte-Torres, Saúl Álvarez-Teijeiro, Sonia Blanco Parajón, Fernando López, Francisco Hermida-Prado, René Rodríguez, Aurora Astudillo, Juana-María García-Pedrero, Iván Fernández-Vega, Juan Pablo Rodrigo, Mónica Álvarez-Fernández

**Affiliations:** Instituto de Investigación Sanitaria del Principado de Asturias (ISPA), Avda. del Hospital Universitario s/n, 33011, Oviedo, Spain; Instituto Universitario de Oncología del Principado de Asturias (IUOPA), University of Oviedo, C/ Julian Clavería s/n 33006 Oviedo, Spain; Spanish Biomedical Research Network in Cancer (CIBERONC), Instituto de Salud Carlos III, Av. Monforte de Lemos, 3-5, 28029, Madrid, Spain; Translational and Clinical Research in Cancer Program, Centro de Investigación del Cáncer, CSIC, Universidad de Salamanca, and FICUS, Campus Miguel de Unamuno s/n, 37007 Salamanca, Spain; Biobanco del Principado de Asturias (BioPA), Avda. del Hospital Universitario s/n, 33011, Oviedo, Spain; Department of Radiation Oncology, Hospital Universitario Central de Asturias, Avenida del Hospital Universitario s/n. 33011, Oviedo, Spain; Department of Otolaryngology, Hospital Universitario Central de Asturias, Avenida del Hospital Universitario s/n. 33011, Oviedo, Spain; Instituto de Biomedicina y Biotecnología de Cantabria (IBBTEC), Universidad de Cantabria-CSIC, Santander, Spain; Service of Pathology, Hospital Universitario Central de Asturias, Avenida del Hospital Universitario s/n. 33011, Oviedo, Spain; Department of Surgery and Medical Surgical Specialties, University of Oviedo, 33006, Oviedo, Spain

**Author notes:** corresponding authors: Juan P. Rodrigo, MD, PhD. Hospital Universitario Central de Asturias Edificio FINBA, Lab ORL Avda. Hospital Universitario s/n, 33011, Oviedo, Spain Mónica Álvarez-Fernández, PhD. Centro de Investigación del Cáncer, CSIC-Universidad de Salamanca-FICUS Campus Miguel de Unamuno s/n, 37007, Salamanca, Spain. These authors contributed equally (Miguel Álvarez-González, Esperanza Pozo-Agundo, Beatriz De Luxán-Delgado).

**Keywords:** patient-derived organoids, head and neck squamous cell carcinoma, laryngeal cancer, pharyngeal cancer, human papillomavirus, cisplatin, radiotherapy, chemotherapy, treatment response, precision oncology

## Abstract

**Background:** Head and neck squamous cell carcinoma (HNSCC) comprises a heterogeneous group of epithelial malignancies associated with poor survival (≈50%), limited therapeutic options, and a lack of predictive biomarkers. Concurrent chemoradiotherapy (CRT) remains the standard treatment for advanced disease; however, many patients fail to respond, develop resistance, or eventually relapse. The development of three-dimensional organoid technology has enabled the generation of patient-derived organoids (PDOs), offering a promising platform for personalized therapeutic testing.

**Methods:** We established a biobank of HNSCC PDOs from fresh laryngeal and pharyngeal tumor samples, including human papillomavirus–positive (HPV+) cases. Organoid formation and expansion rates were analyzed in relation to clinical parameters. Selected representative PDOs were histologically and molecularly characterized. Additionally, several models were exposed to cisplatin and radiation to evaluate treatment response, and a subset was assessed for tumorigenicity in subcutaneous mouse models.

**Results:** Fifty-seven PDO models were successfully established, long-term expanded, and cryopreserved. Prior chemotherapy and/or radiotherapy was identified as an independent negative predictor of organoid outgrowth and expansion capacity compared with treatment-naïve samples. Histological features, including differentiation grade and immunohistochemical markers, were largely preserved and strongly correlated with the original tumors. PDOs displayed heterogeneous responses to cisplatin and radiotherapy, with HPV-positive models showing greater sensitivity, consistent with clinical observations. Global transcriptomic profiling revealed molecular subtypes concordant with established HNSCC classifications and suggested an additional subtype characterized by low MYC and mTORC1 transcriptional activity.

**Conclusion:** HNSCC PDOs faithfully recapitulate tumor histology and molecular diversity, providing a robust platform to investigate tumor biology and therapeutic response.

## INTRODUCTION

Head and neck squamous cell carcinoma (HNSCC), arising from the mucosa of the oral cavity, pharynx, or larynx, remains a malignancy with high mortality and limited therapeutic options. Tobacco and alcohol consumption, as well as human papillomavirus (HPV) infection—particularly in oropharyngeal tumors—constitute the primary risk factors. Although patients diagnosed at early-stage achieve 5-year overall survival (OS) rates of 70–90%, more than 60% present with advanced tumors, for which OS remains poor, especially in HPV-negative cases, with minimal improvement over the past three decades [1]. Standard treatment for advanced disease relies on surgery and/or radiotherapy (RT) often combined with cisplatin-based chemotherapy (CT). In addition, the only approved targeted therapy for HNSCC is the anti-EGFR monoclonal antibody cetuximab [2]; however, its clinical efficacy is limited, with durable responses only observed in a small subset of patients. More recently, immune-checkpoint inhibitors targeting PD-1, such as pembrolizumab and nivolumab, have expanded the therapeutic landscape for HNSCC [3]. Despite these advances, clinical responses remain modest, and a major unmet need is the lack of robust predictive biomarkers capable of anticipating therapeutic response and guiding treatment decisions.

Conventional HNSCC cell lines have been widely used in translational research; however, their clinical relevance is often limited, likely due to their two-dimensional nature and the acquisition of genetic and phenotypic alterations during long-term in vitro culture. In contrast, murine models better recapitulate tumor physiology but present important limitations, including high cost, long experimental timelines and limited suitability for high-throughput drug screening. These challenges have promoted the development of New Approach Methodologies (NAMs) aiming at reducing animal use while more faithfully modeling human disease. Among these, three-dimensional organoid models and, in particular, patient-derived organoids (PDOs) have emerged as powerful preclinical platforms for translational research [4]. Tumor organoids, or tumoroids, are self-organizing three-dimensional structures derived from patient tumor stem cells that can be expanded long-term and cryopreserved *in vitro*. Importantly, they retain the multi-omic features of their tumor of origin, making them highly attractive for functional precision oncology [5].

Protocols for the establishment of HNSCC PDOs were first described in 2019 by Driehuis *et al* [6]. That initial biobank of 31 HNSCC-derived organoids was subsequently expanded to over 100 models, predominantly derived from oral cavity tumors [5]. Since then, several recent independent studies have reported the generation and characterization of additional HNSCC PDO collections, most of which are similarly enriched in oral cavity-derived tumors [7-9]. A relatively underexplored aspect of these models is their ability to recapitulate the differentiation program of the stratified epithelium of the upper aerodigestive tract and, consequently, to reflect the histological grade of the tumors from which they originate.

Here we report the establishment of a large robust collection of pharyngeal and laryngeal HNSCC PDOs, including HPV-positive cases, that faithfully recapitulate distinct tumor differentiation states. These models also reproduce differential responses to standard-of-care therapies, with HPV-positive PDOs showing the most favorable responses. Notably, a subset of PDOs does not conform to previously defined molecular subtypes, suggesting the existence of at least one potential new subtype. Intriguingly this subgroup exhibits the highest cisplatin resistance in vitro and is associated with increased rates of tumor recurrence.

## MATERIALS AND METHODS

### Collection of human samples and clinical data

Fresh tumor samples were obtained between 2020 and 2024 from patients with HNSCC (including laryngeal, hypopharyngeal, and HPV-positive or -negative oropharyngeal tumors) recruited by the Department of Otorhinolaryngology at the Hospital Universitario Central de Asturias (HUCA), who underwent surgical treatment or needed a biopsy for diagnostics. This study was approved by the Regional Ethics Committee of the Principality of Asturias (CEImPA) (date of approval 6 April 2020; approval number 2020.064, for the project INVES19001ALVA and date of approval 25 January 2021; approval number 2021.002, for the project PID2020-117236RB-I00). Written informed consent was obtained from all patients. All tissue samples were collected via the Biobank of the Principality of Asturias (BioPA, National Registry of Biobanks B.0000827). Clinical patient-related data were annotated. Patient age was defined as the age at the time of sample collection.

### Tissue processing for organoid establishment

Organoids were established and maintained as described by Driehuis *et al*. [6] with minor modifications. Tissue specimens were collected in basal media, consisting of Advanced DMEM/F12 (AdDMEM/F12: Gibco 12634028), supplemented with 1x GlutaMAX (Thermofisher; Gibco, cat # 35050061), penicillin-streptomycin (Biowest; cat #L0022-100) and 10 mM HEPES (Gibco, cat # 15630056). Tissue samples were minced into small fragments, washed in basal medium and digested for 30-60 min at 37 °C in 5 mL of 0.125% Trypsin (Sigma, cat # T1426) prepared in basal medium. Mechanical dissociation was enhanced by pipetting every 10 min. Digestion was monitored at the microscope and stopped by adding 10 ml of cold basal medium. The suspension was filtered through a 100 µm Easy-Strainer filter (Corning; #431752) and centrifuged at 300 x g for 5 min at 4 °C. When needed cell pellet was incubated with ACK lysing buffer (Gibco, cat # A1049201) for 5-10 min at room temperature (RT) to remove red blood cells, washed with basal medium and centrifuged at 300 x g for 5 min at 4 °C. Cell pellet was gently resuspended in ice-cold 80-90% Matrigel (Corning; #356231) in basal medium and plated as ∼10µL domes (four domes per well) in pre-warmed untreated 24-well culture plates (Nunc #144530, ThermoScientific) and left in a 37 °C/ 5% CO_2_ incubator upside-down for 20-30 min until polymerization. Then, 500 µL of pre-warmed PDO expansion medium was added, supplemented with 10 µM Y-27632 (Tocris #1254); 0.5 µg/mL caspofungin (Sigma #SML0425); and 100 µg/mL Primocin (InvivoGen #ant-pm-05). Cultures were incubated at 37 °C in a humidified atmosphere with 5% CO₂. Two types of expansion culture media were used: HN (M0) medium, as described by Driehuis *et al*. [6] or cervical SCC medium (M7), as described by Lohmussaar *et al*. [10], with slight modifications.

HN (M0) expansion medium consisted of basal medium supplemented with B27 (Gibco #17504044), 10 mM nicotinamide (Sigma #N0636), 1.25 mM N-acetylcysteine (Sigma #A9165), 500 nM A83-01 TGFbR-ALK5 inhibitor (Tocris #2939), 1 µM PGE-2 Prostaglandin E2 (Tocris #2296), 0.3 µM CHIR99021 GSK3 inhibitor (SIGMA #SML1046), 1 µM forskolin (Tocris #1099), 50 ng/mL hEGF (Preprotech #AF-100-15), 10 ng/mL hFGF10 (Preprotech #AF-100-26), 5 ng/mL hFGF2 (Preprotech #AF-100-18B), 100 ng/mL noggin (Preprotech #120-10C), and 200 ng/mL R-spondin (hRspo) (Preprotech #120-38).

M7 expansion medium consisted of basal medium supplemented with B27 (Gibco #17504044), 10 mM nicotinamide (Sigma #N0636), 1.25 mM N-acetylcysteine (Sigma #A9165), 500 nM A83-01 TGFbR-ALK5 inhibitor (Tocris #2939), 1µM SB202190 (Sigma Aldrich # S7067), 10 µM forskolin (Tocris #1099), 100 ng/mL hFGF10 (Preprotech #AF-100-26), 25 ng/mL hFGF7 (Preprotech #100-19), 100 ng/mL noggin (Preprotech #120-10C), and 200 ng/mL R-spondin (hRspo) (Preprotech #120-38).

All cultures were initially grown in HN expansion medium. M7 medium was incorporated into the study from 2023 onward following a published report demonstrating improved growth and biobanking efficiency [11]. An overview of the culture conditions used for each sample is provided in **Table S1**. Medium was changed every 2-3 days and organoids were passaged approximately 7-14 days after plating, depending on their growth rate.

### Organoid culturing and passaging

For passaging, Matrigel droplets containing organoids were mechanically disrupted by scraping and resuspending the well content using a P1000 pipette. The suspension was transferred to a 15 mL Falcon tube and brought up to 10 mL with basal medium. Samples were centrifugated at 300 x g, for 5 min, at room temperature (RT). The supernatant was discarded and the pellet was resuspended and incubated for 5-15 min at 37 °C in 1-3 mL of TryplE Express 1X (Gibco; #12605010). During enzymatic dissociation, organoids were mechanically sheared every 2–3 min by pipetting with a P1000 pipette fitted with an additional P10 tip to increase shear stress. Digestion was closely monitored microscopically and stopped once organoids were dissociated into single cells or small clusters. Tubes were then filled up to 15 mL with basal medium, centrifuged at 300 x g for 5 min at RT, and kept on ice until the supernatant was removed and the pellet resuspended in 80–90% Matrigel diluted in basal medium.

As an optional step, prior to passaging, 1 mg/mL of Dispase II (Sigma; #D4693-1G) was added to each well and incubated at 37 °C for 30 min to facilitate matrix dissociation before organoid collection.

Organoid density and appropriate split ratios were assessed microscopically before passaging and confirmed by pellet size after centrifugation. As described for primary tumor seeding, organoids were embedded in four ∼10 µL Matrigel domes per well in pre-warmed, untreated plates, which were then inverted and incubated at 37 °C for 20–30 min to allow Matrigel polymerization. Subsequently, 500 µL of pre-warmed expansion medium supplemented with 10 µM Y-27632 was added per well. Expansion medium was replaced every 2–3 days, and Y-27632 was omitted after the first medium change.

### Freezing and thawing organoids

Organoids were passaged and expanded, and day 2–3 Matrigel domes were disrupted using 1 mg/mL Dispase II, as described above. Following Dispase incubation, organoids were collected diluted in basal medium, and centrifuged. The supernatant was discarded, and the organoid pellet was washed with basal medium and centrifuged again. Pellet was gently resuspended in cold Recovery Cell Freezing Medium (Gibco, #12648010) by dropwise addition (500 µL per cryovial, corresponding to material from approximately 2–4 wells).

For recovery, cryovials were rapidly thawed in a 37 °C water bath and the contents were immediately transferred to a pre-warmed tube containing 10 mL of basal medium. Samples were centrifuged, placed on ice, and the supernatant was completely removed. Pellets were resuspended in Matrigel (50 µL per well) and plated as ∼10 µL domes in pre-warmed 24-well plates. After Matrigel solidification, cultures were overlaid with expansion medium supplemented with Y-27632.

### Histology and immunohistochemistry

Organoids were harvested 7-10 days after passaging following Matrigel dissociation with Dispase II, as described above. The organoid pellet was washed with PBS, fixed in 4% formalin for at least 4 hours or overnight at RT, and stored in PBS at 4°C until processing. The pellet was stained with hematoxylin, washed with PBS, and placed between two biopsy pads inside a cassette for paraffin embedding. Paraffin blocks containing organoids or tumor tissue fragments were sectioned onto glass slides and subjected to hematoxylin and eosin (H&E) and immunohistochemical (IHC) staining.

The following primary antibodies were used: monoclonal mouse anti-human Ki-67 antibody (clone MBI-1; Dako, #GA626), monoclonal mouse anti-CDKN2A/p16INK4a antibody (clone JC8; Santa Cruz, #sc-56330; 1:50 dilution), monoclonal mouse anti-human p63 (Clone DAK-p63; Dako/Agilent #IR662) and mouse monoclonal anti-keratin K13 (Ks13.1; Progen #61007; 1:50 dilution) Detection was performed using the Dako EnVision Flex. Visualization System (Dako Autostainer).

Slides were scanned using a NanoZoomer-SQ Digital Slide Scanner C13140-01 (Hamamatsu) and analyzed with NDP.scan and NDP.View software (version 1.0.9).

H&E and IHC sections from tumors and their corresponding PDOs were evaluated by a pathologist to assess the degree of histopathological correlation between the primary tumors and their matched PDOs.

### Cisplatin and radiotherapy treatments

Organoids were collected 2–3 days after passaging following Matrigel dissociation with Dispase II, as described above. Samples were filtered through a 70 µm cell strainer (Falcon, #352350), counted, and dispensed into pre-warmed 96-well untreated plates (Nunc, Thermo Fisher Scientific, #236105). A total of 2,000 organoids were seeded per well in a 5 µl dome of 95% Matrigel in basal medium, with at least three technical replicates per condition. After polymerization, 150 µL of pre-warmed PDO expansion medium was added.

Two to three days later (i.e., organoid day 5), cultures were treated with increasing concentrations of cisplatin (0, 0.6, 1.25, 2.5, 5, 10, and 20 µM) in expansion medium for 5 days. Cell viability was assessed by measuring intracellular ATP levels using the CellTiter-Glo assay 3D (Promega, #G9681). Briefly, culture medium was replaced with 100 µL of basal medium per well, followed by the addition of 100 µL of CellTiter-Glo 3D reagent. Plates were incubated at room temperature, covered with foil, on a shaker for 30 min to ensure complete organoid lysis. Luminescence was measured using a Synergy HT plate reader (BioTek, Winooski, VT, USA).

For radiotherapy experiments, organoids were prepared as described above. On day 5, organoids were irradiated with a single dose of 4 or 8 Gy using a square PMMA phantom as a culture plate adaptor for a linear accelerator (TrueBeam, Varian) at the Radiation Oncology Unit (HUCA). A non-irradiated plate (0 Gy) was included as a negative control. Expansion medium was replaced immediately after irradiation, and cell viability was assessed 5 days later using the CellTiter-Glo 3D assay as described above.

For chemoradiotherapy experiments, cisplatin treatment was administered the day after irradiation. Culture medium was removed and organoids were treated with a single dose of cisplatin (1 or 5 µM). Viability was measured 4 days after cisplatin treatment using the same ATP-based assay described above.

A CellTiter-Glo 3D assay was also performed on the day of seeding to establish baseline viability and estimate growth rates throughout the experiment.

### RNA isolation

Organoids were collected 7–10 days after seeding following Matrigel disruption with Dispase II, as previously described. The resulting pellet was stored at −80 °C until processing. Total RNA was extracted using the GeneJET RNA Purification Kit (Thermo Fisher Scientific, #K0731) according to the manufacturer’s instructions.

### RNA sequencing analysis

RNA sequencing was performed by Novogene (Cambridge, UK). mRNA libraries were prepared by poly(A) enrichment and sequenced on an Illumina NovaSeq X Plus platform (paired end, 150 bp). An average of 20 million reads per sample was obtained (∼6 Gb per sample).

Raw sequencing data were subjected to quality control, and adapter sequences and low-quality paired-end reads were removed using fastp (v0.23.4). Preprocessed reads were pseudo-aligned to the GRCh38 human genome and quantified at the transcript level using the mapping-based mode of Salmon (v1.10.3). The Salmon index was generated from a gentrome constructed from the GENCODE v46 transcriptome and the GRCh38.p14 primary genome assembly.

Gene expression analyses were conducted in R (v4.4.1) using Bioconductor packages (v3.19). Transcript-level estimates were summarized to gene-level counts for exploratory and differential expression analyses using tximeta (v1.22.1). For visualization, raw count matrices were variance-stabilizing transformed (VST) using DESeq2 (v1.44.0). Lowly expressed genes (<10 counts in fewer than three samples) were filtered out to reduce the multiple testing burden.

Differential expression analysis was performed using the negative binomial generalized linear model framework implemented in DESeq2. P-values were adjusted for multiple testing using the Benjamini–Hochberg method. Differentially expressed genes (DEGs) were defined as those with a false discovery rate (FDR) < 0.05 and an absolute log_2_ fold change > 1.

Sample clustering was performed by grouping samples with a Spearman correlation coefficient ≥ 0.9, and the optimal number of clusters was determined using the sum of squared errors (SSE) method. Gene annotation was performed using the org.Hs.eg.db package (v3.19.1) to map Ensembl Gene IDs to Entrez Gene IDs, gene symbols, biotypes, and gene descriptions.

Pathway enrichment analysis between conditions was conducted using clusterProfiler (v.4.12.6) with gene sets from the Molecular Signatures Database (MSigDB) Hallmark collection, accessed through msigdbr package (v.7.5.1).

Transcription factor activity was inferred using decoupleR (v.2.10.0) with the collecTRI collection and the univariate linear model (ULM) method.

### Mouse xenografts

All mouse experiments were performed in accordance with the institutional guidelines of the University of Oviedo and were approved by the Animal Research Ethics Committee of the University of Oviedo prior to study initiation (approval date: 1 August 2019, approval number PROAE 46/2019; approval date: 1 April 2024, approval number PROAE 03/2024).

Female Athymic Nude-*Fox1nu* (Hsd) mice and NOD.CB17-*Prkdc*^scid^ *Il2rg*^tm1^/BcgenHsd (B-NDG) mice aged 5–6 weeks were purchased from Envigo (Inotiv, Inc).

Prior to transplantation, PDO organoid lines were cultured in Matrigel droplets as described above but scaled-up to 6-well plates. On the day of transplantation, day-5 organoids were collected by Dispase II–mediated Matrigel disruption, as previously described. A small fraction of the sample was dissociated into single cells to estimate the cellular density of the PDO preparation. Mice were subcutaneously inoculated in the flanks with day-5 formed organoids (approximately 1.5 × 10⁶ cells) resuspended in 100 µL of expansion medium mixed 1:1 with Matrigel.

Tumor size was measured using a caliper once or twice per week, and tumor volume was calculated using the formula V = π (D × d²)/6, where D represents the maximum diameter and d the minimum diameter.

Animals were euthanized by cervical dislocation 70 days after transplantation. Tumors that had successfully developed were excised from the flanks, fixed in formalin, embedded in paraffin, and processed for H&E staining.

### Statistical analysis

Associations with clinicopathological features were evaluated using SPSS software (IBM). Chi-square (χ²) and Fisher’s exact tests were applied to compare categorical variables. Multivariate logistic regression was performed using the backward likelihood ratio (LR) method to determine the independent effect of each parameter. Odds ratios (ORs), 95% confidence intervals (CIs), and *p* values were reported. Comparisons between two groups were performed using an unpaired Student’s t-test, whereas comparisons among more than two groups were conducted using one-way ANOVA, using Prism version 10.6.1 (GraphPad). All tests were two-sided, and *p* values ≤ 0.05 were considered statistically significant.

## RESULTS

### Establishment of a collection of HNSCC PDOs

From May 2020 to December 2024, a total of 180 samples were processed from 167 patients with pharyngeal or laryngeal squamous cell carcinomas spanning various disease stages. Detailed patient characteristics are presented in **Table 1**.

**Table 1.** Associations between organoid establishment and clinicopathological factors.

| Feature | N<br>samples<br>(%) | Organoid formation |  | Organoid expansion |  |
| --- | --- | --- | --- | --- | --- |
|  |  | Yes<br>(derivation rate) | p value | Yes<br>(freezing rate) | p value |
| **- Sex** |  |  |  |  |  |
| Male | 169 (94) | 116 (69) | 1<sup>†</sup> | 55 (33) | 0.506<sup>†</sup> |
| Female | 11 (6) | 8 (73) |  | 2 (18) |  |
| **- Age** |  |  |  |  |  |
| Below median ($\leq 68.4$) | | 58 (64) | 0.26<sup>†</sup> | 29 (32) | 1<sup>†</sup> |
| Above median ($> 68.4$) | | 66 (73) | | 28 (31) | |
| **- T classification (n=151)** |  |  |  |  |  |
| T1-T2 | 68 (45) | 41 (60) | 0.226<sup>#</sup> | 18 (26) | 0.345<sup>#</sup> |
| T3 | 49 (32.5) | 36 (73) |  | 19 (39) |  |
| T4 | 34 (22.5) | 25 (73) |  | 12 (35) |  |
| **- N classification (n=141)** |  |  |  |  |  |
| N0 | 77 (55) | 50 (65) | 0.468<sup>†</sup> | 26 (34) | 1<sup>†</sup> |
| N1-3 | 64 (45) | 46 (72) |  | 21 (33) |  |
| **- Disease stage (n=127)** |  |  |  |  |  |
| I-II | 44 (34) | 29 (66) | 0.390<sup>#</sup> | 15 (34) | 0.821<sup>#</sup> |
| III | 25 (20) | 17 (68) |  | 10 (40) |  |
| IV | 58 (46) | 45 (77) |  | 23 (40) |  |
| **- Pathological grade (n=160)** |  |  |  |  |  |
| G1 | 49 (31) | 32 (65) | 0.650<sup>#</sup> | 15 (31) | 0.650<sup>#</sup> |
| G2 | 79 (49) | 55 (70) |  | 27 (34) |  |
| G3 | 32 (20) | 24 (75) |  | 13 (41) |  |
| **- Anatomic site** |  |  |  |  |  |
| Oropharynx | 46 (26) | 34 (74) | **0.022<sup>#</sup>** | 14 (30) | 0.660<sup>#</sup> |
| Hypopharynx | 21 (12) | 9 (43) |  | 5 (24) |  |
| Larynx | 113 (63) | 81 (72) |  | 38 (34) |  |
| **- HPV status (n=46)‡** |  |  |  |  |  |
| Negative | 30 (65) | 18 (60) | **0.004<sup>†</sup>** | 6 (20) | **0.048<sup>†</sup>** |
| Positive (p16+) | 16 (35) | 16 (100) |  | 8 (50) |  |
| **- Treatment** |  |  |  |  |  |
| Treatment naïve | 131 (73) | 100 (76) | **<0.001<sup>†</sup>** | 44 (34) | **0.029<sup>†</sup>** |
| Surgery | 19 (10) | 14 (74) |  | 9 (47) |  |
| RT/CRT/CT | 30 (17) | 10 (33) |  | 4 (13) |  |
| **- Sample type** |  |  |  |  |  |
| Surgical specimen | 150 (83) | 102 (68) | 0.668<sup>†</sup> | 49 (33) | 0.668<sup>†</sup> |
| Office biopsy | 30 (17) | 22 (73) |  | 8 (27) |  |
| **Total Cases** | **180** | **124 (69)** |  | **57 (32)** |  |

Organoid formation was observed in 69% of samples, and efficient expansion with successful cryopreservation was achieved in 32%. These rates increased to 75% and 35%, respectively, when only considering samples in which epithelial cells were visible during digestion. No epithelial cells were detected after digestion in 17 of these samples (9%).

The median age of patients whose tumor samples were able to form organoids was 69, comparable to 67.8 years in samples that did not form organoids. Similarly, no significant differences were found in median age between successfully expanded and cryopreserved organoids (66.8) and those unable to expand (68.7 years). No significant differences were found for other clinicopathological variables such as tumor size, lymph node infiltration, disease stage or differentiation grade. No differences were detected based on the type of sample (surgical vs. office). However, a significant difference in organoid formation efficiency was observed depending on the anatomical site, with hypopharyngeal tumors being markedly less efficient (43%) compared to laryngeal or oropharyngeal tumors (approximately 70%). Among the latter tumors, HPV-positive cases were the most efficient in terms of both organoid formation (100% vs 60%) and expansion (50% vs 20%) compared with HPV-negative tumors. Notably, the strongest association was observed with pre-treatment status. Both organoid formation (33% vs 74-76%, *p* < 0.001) and expansion (13% vs 34-47%, *p* = 0.029) were significantly reduced when tissue samples were obtained from patients previously treated with CT, RT or CRT compared to treatment-naïve patients or those pretreated by surgery alone **(Table 1)**.

Next, a binary multivariable logistic regression using the backward likelihood ratio (LR) method was conducted to identify independent predictors of organoid formation and expansion. Here, large tumor size emerged as an independent factor strongly associated with organoid formation (T3, OR = 2.613, 95%CI [1.011-6.752], *p* = 0.047; T4, OR =3 .835, 95%CI [1.121-13.125], *p* = 0.032), while HPV positivity significantly increased the probability of organoid expansion (OR = 3.547, 95%CI [1.047-6.752], *p* = 0.042) **(Table 2)**. Interestingly, pre-treatment with CT and/or RT remained as independent significant predictor of poor organoid formation (OR=0.098, 95%CI [0.027,

**Table 2.** Multivariate analysis of organoid establishment and clinicopathological parameters.

| Variable | Organoid formation |  | Organoid expansion |  |
| --- | --- | --- | --- | --- |
|  | OR (95%CI) | p value | OR (95%CI) | p value |
| **Sex** (male vs female) | 1.660 (0.258-10.693) | 0.594 | 0.432 (0.072-2.598) | 0.359 |
| **Age** (below vs above median) | 0.982 (0.388-2.484) | 0.969 | 0.620 (0.277-1.387) | 0.245 |
| **T** (*T1-T2 reference*) |  |  |  |  |
| T3 | **2.613 (1.011-6.752)** | **0.047** | 2.339 (0.945-5.788) | 0.066 |
| T4 | **3.835 (1.121-13.125)** | **0.032** | 1.872 (0.651-5.383) | 0.245 |
| **N** (N0 vs N+) | 0.776 (0.249-2.423) | 0.663 | 0.921 (0.388-2.185) | 0.852 |
| **Grade** (*reference G1*) |  |  |  |  |
| G2 | 2.468 (0.844-7.218) | 0.099 | 1.035 (0.381-2.815) | 0.946 |
| G3 | 2.243 (0.604-8.332) | 0.214 | 1.128 (0.338-3.761) | 0.845 |
| **Site** (*reference: oropharynx*) |  |  |  |  |
| Hypopharynx | 0.277 (0.051-1.522) | 0.140 | 0.851 (1.145-4.981) | 0.858 |
| Larynx | 0.717 (0.183-2.808) | 0.633 | 0.891 (0.215-3.694) | 0.874 |
| **HPV** (negative or not determined vs positive) | 1.6e9 (N/A extreme value) | 0.998 | **3.547 (1.047-12.011)** | **0.042** |
| **Treatment** (*reference naïve*) |  |  |  |  |
| surgery | 1.386 (0.334-5.743) | 0.653 | 2.626 (0.831-8.298) | 0.100 |
| RT/CRT/CT | **0.098 (0.027-0.366)** | **0.001** | 0.221 (0.047-1.034) | 0.055 |
| **Sample** (surgical vs office) | 0.404 (0.120-1.361) | 0.143 | 0.509 (0.169-1.537) | 0.231 |

OR, odds ratio; CI, confidence interval

0.366], *p* = 0.001) and was negatively associated with organoid expansion, approaching statistical significance (OR = 0.221, 95%CI [0.047-1.034], *p* = 0.055) **(Table 2)**. These data strongly confirm that prior treatment with CT and/or RT negatively impacts the ability to establish and expand organoid cultures from patient tumor samples.

Successfully expanded and frozen organoids (n=57) were enriched in laryngeal tumors followed by oropharyngeal and hypopharyngeal ones **(Figure 1a and Supplementary Table S1)** and in moderately differentiated G2 tumors **(Figure 1b)**. Recovery after thawing was evaluated in 39 models: 23 (59%) were successfully recovered and further expanded, 6 (15%) were lost upon thawing, and 10 (25%) showed uncertain and/or low recovery efficiency, requiring additional testing **(Supplementary Table S1)**.

**Figure 1.**
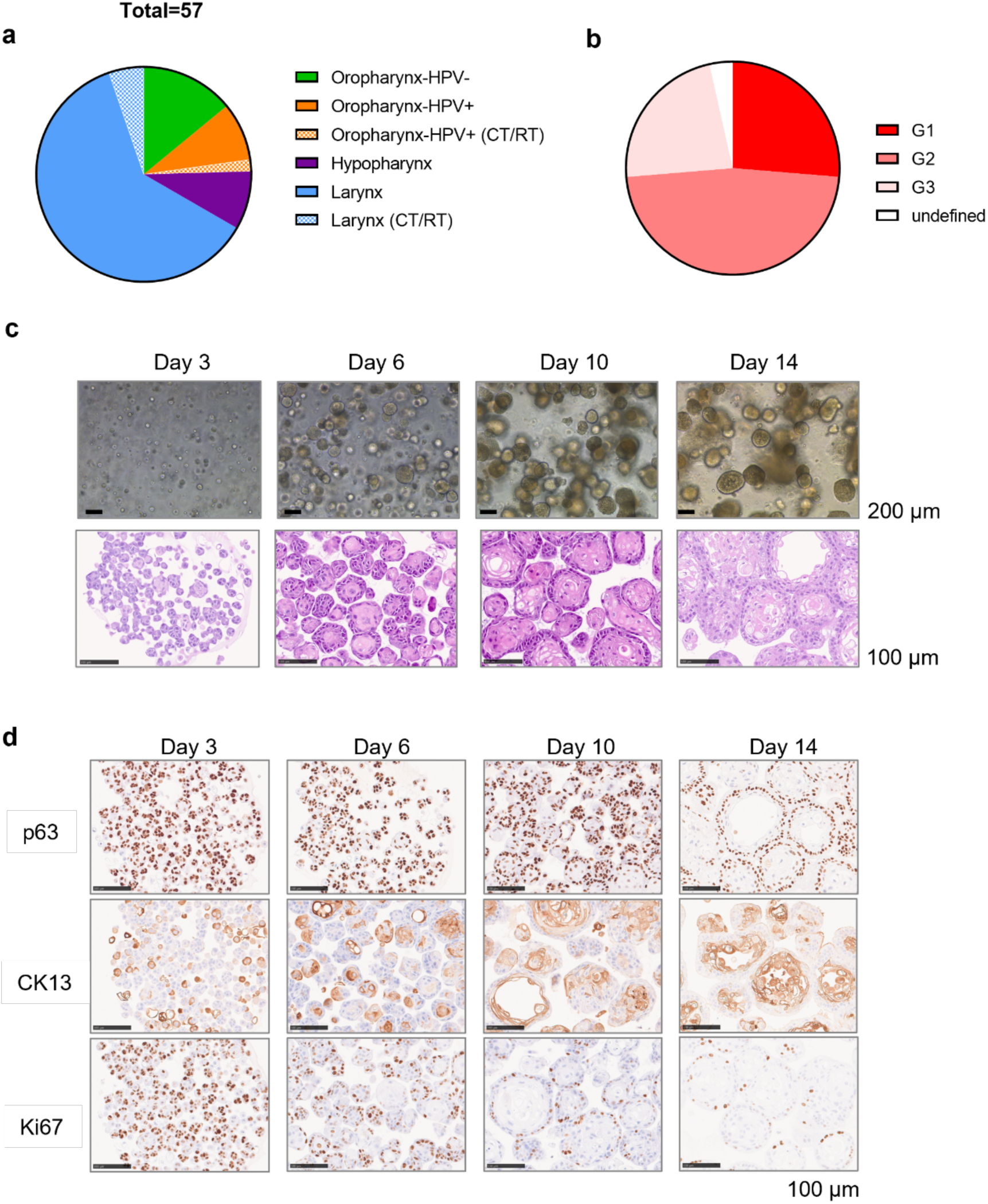
Generation and characterization of a patient-derived organoid (PDO) collection from pharyngeal and laryngeal squamous cell carcinomas. **a)** Composition of the HNSCC PDO collection according to the anatomical origin of the tumors from which organoids were established. CT/RT in brackets indicate organoids derived from patients previously treated with chemotherapy (CT) and/or radiotherapy (RT). HPV, human papillomavirus. **b)** Distribution of PDO models according to the histological grade of the corresponding primary tumors (see also **Table S1**). **c)** Representative bright-field images of a PDO culture at the indicated days after passaging, alongside with matched hematoxylin and eosin (H&E) staining of formalin-fixed, paraffin-embedded (FFPE) organoids at the same time points after splitting. Scale bars, 200 µm (*top panel*); 100 µm (*bottom panel*) **d)** Immunohistochemical analysis of the basal marker p63 (*top panel*), the differentiation marker CK13 (*middle panel*) and the proliferation marker Ki-67 (*bottom panel*) in a representative PDO model at different time points after passaging. Scale bars: 100 µm.

In contrast to other epithelial organoids, HNSCC PDOs are mostly compact, reflecting their origin from a stratified epithelium **(Figure 1c)**. Notably, organoid establishing and expansion recapitulate the differentiation process of a stratified epithelium, starting with organoids composed almost entirely of undifferentiated basal cells (marked by the basal marker p63) in the early days after passage, and progressively increasing in differentiated (cytokeratin 13, CK13-positive) cells as the organoids mature and grow in size, particularly in those exhibiting moderate or well-differentiation patterns **(Figure 1d)**.

In addition, we compared the success rate of organoid establishment in the original head and neck expansion medium (HN) described by Driehuis *et al*. in 2019 [6] with the M7 medium described for cervical organoids [10], which has recently been shown to be more efficient for HNSCC PDOs [11]. Consistent with those data, we also found higher efficiency in M7 medium compared to the traditional HN (M0) medium (**Figure S1).** Of the 13 samples plated in both media, most of them grew better in M7 or showed no differences, with only one sample performing better in HN (M0) **(Figure S1a)**. In general, organoids expanded in M7 medium showed a higher number and smaller size, with no noticeable morphological differences **(Figure S1b and b)**. Therefore, we decided to switch to M7 medium as the primary derivation medium. Interestingly, when analyzing separately the subgroup of organoids derived in HN medium (from 2020 to July 2023) and those established in M7 medium (from August 2023 onwards), we did not find major differences in their association with clinical features **(Supplementary Tables S2 and S3, respectively)** compared to the entire cohort **(Table 1)**. Importantly, lower efficiency in both organoid formation and expansion was again confirmed in organoids from CT/RT pre-treated tumors established and cultured either in HN or M7 medium.

### HNSCC PDOs recapitulate the differentiation grade of their tumor of origin

To assess whether the histological features of the tumors of origin were preserved in the derived organoid models, we selected 13 organoid cultures representing different anatomical sites and histological grades and compared them with their corresponding primary tumors. We observed a strong concordance in differentiation grade and IHC marker expression in 10 of the 13 paired samples (77%; Spearman r = 0.80, *p* = 0.002) **(Figure 2; Supplementary Table S4)**.

**Figure 2.**
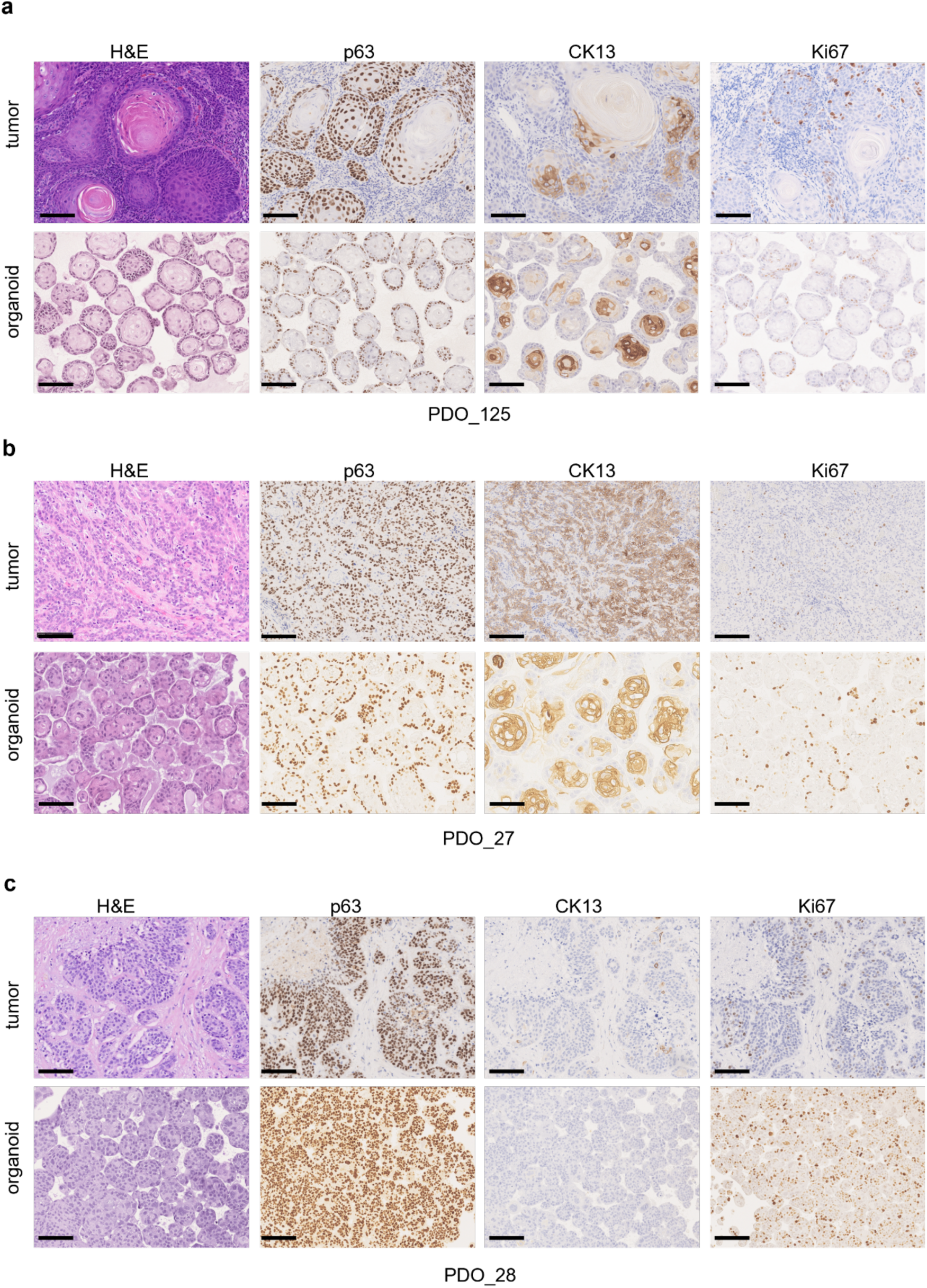
Morphological evaluation of HNSCC tumors and matched PDOs. Representative images of H&E staining and immunohistochemical detection of p63 (basal marker), CK13 (squamous differentiation marker) and Ki-67 (proliferation marker) stainings in three HNSCC tumors and their matched PDOs, illustrating different histological grades: **a)** G1, well-differentiated (PDO_125, *top panel*); **b)** G2, moderately differentiated (PDO_27, *middle panel*); and **c**) G3, poorly differentiated (PDO_28, *bottom panel*). Scale bars: 100 µm.

Organoids derived from HPV-positive tumors also recapitulated key histological characteristics of their primary counterparts, including the presence of koilocytes indicative of HPV infection, high p16 expression, and generally poorer differentiation grade **(Figure 3)**.

**Figure 3.**
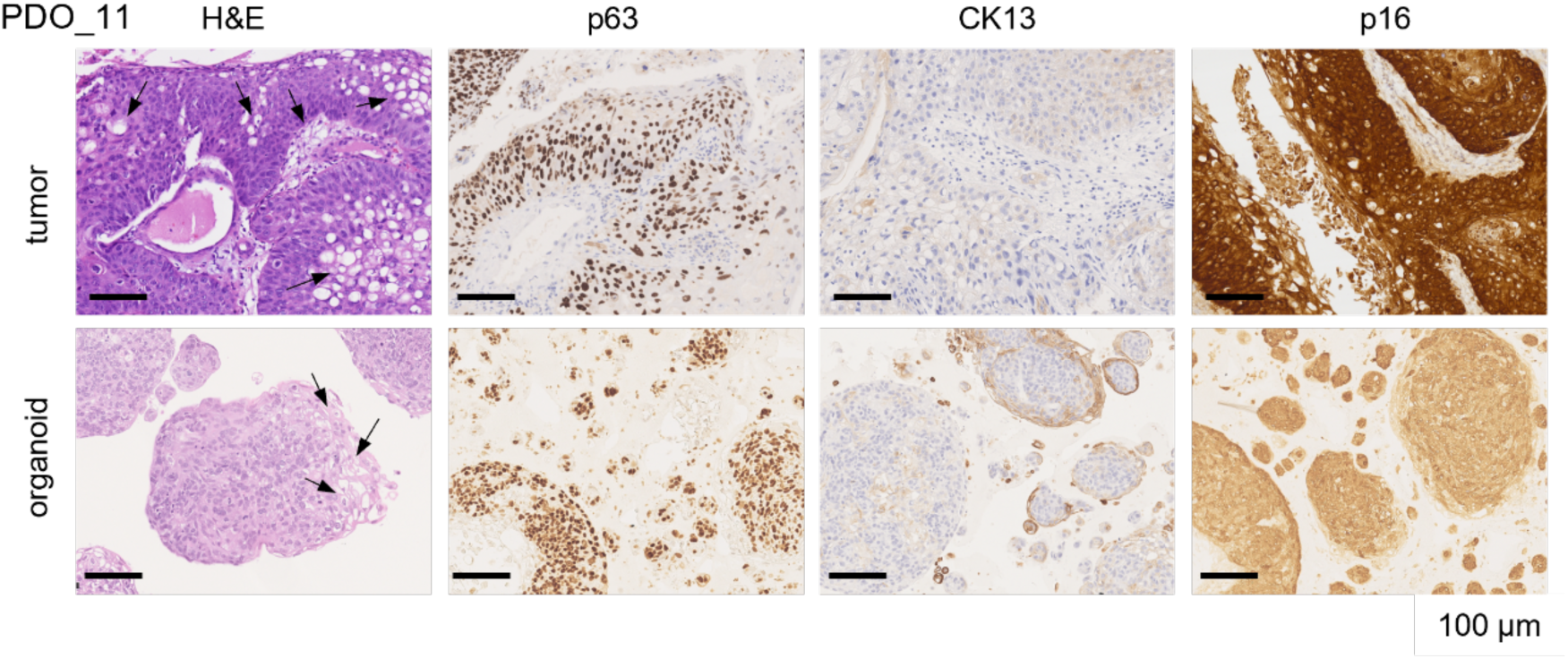
HNSCC PDOs recapitulate morphological features of HPV-positive tumors. Histological and immunohistochemical analysis of a representative HPV-positive oropharyngeal squamous cell carcinoma and its matched PDO model. Staining for p63 (basal marker), CK13 (squamous differentiation marker) and p16 (surrogate marker of HPV infection) is shown. Arrows mark koilocytes. Scale bars: 100 µm.

Among the remaining three cases, one model (PDO_225) showed poor concordance: the organoid was classified as moderately differentiated, whereas the tumor of origin was a poorly differentiated laryngeal squamous cell carcinoma (SCC), despite exhibiting high CK13 expression. The other two cases showed partial concordance. In PDO_108, the organoid displayed a higher degree of differentiation compared with its tumor of origin, which was a highly heterogeneous laryngeal SCC containing both differentiated and undifferentiated areas. PDO_210 showed marked intra-culture heterogeneity, with a mixture of moderately and well-differentiated organoids, whereas the originating oropharyngeal tumor was moderately to poorly differentiated **(Supplementary Figure S2)**.

### HNSCC PDOs show differential responses to chemo- and radiotherapy

To evaluate organoid sensitivity to standard-of-care treatments for HNSCC, established PDOs were exposed to increasing doses of cisplatin or ionizing radiation (IR), and organoid viability was assessed using an ATP-based luminescence assay **(Figure 4a)**. PDOs displayed heterogeneous responses to cisplatin. HPV-positive PDOs were the most sensitive, with IC₅₀ values ranging from 1–5 µM. In contrast, HPV-negative oropharyngeal PDOs showed lower sensitivity (IC₅₀ >10 µM), whereas laryngeal PDOs exhibited broader variability, with IC₅₀ values spanning 2–20 µM **(Figure 4b)**.

**Figure 4.**
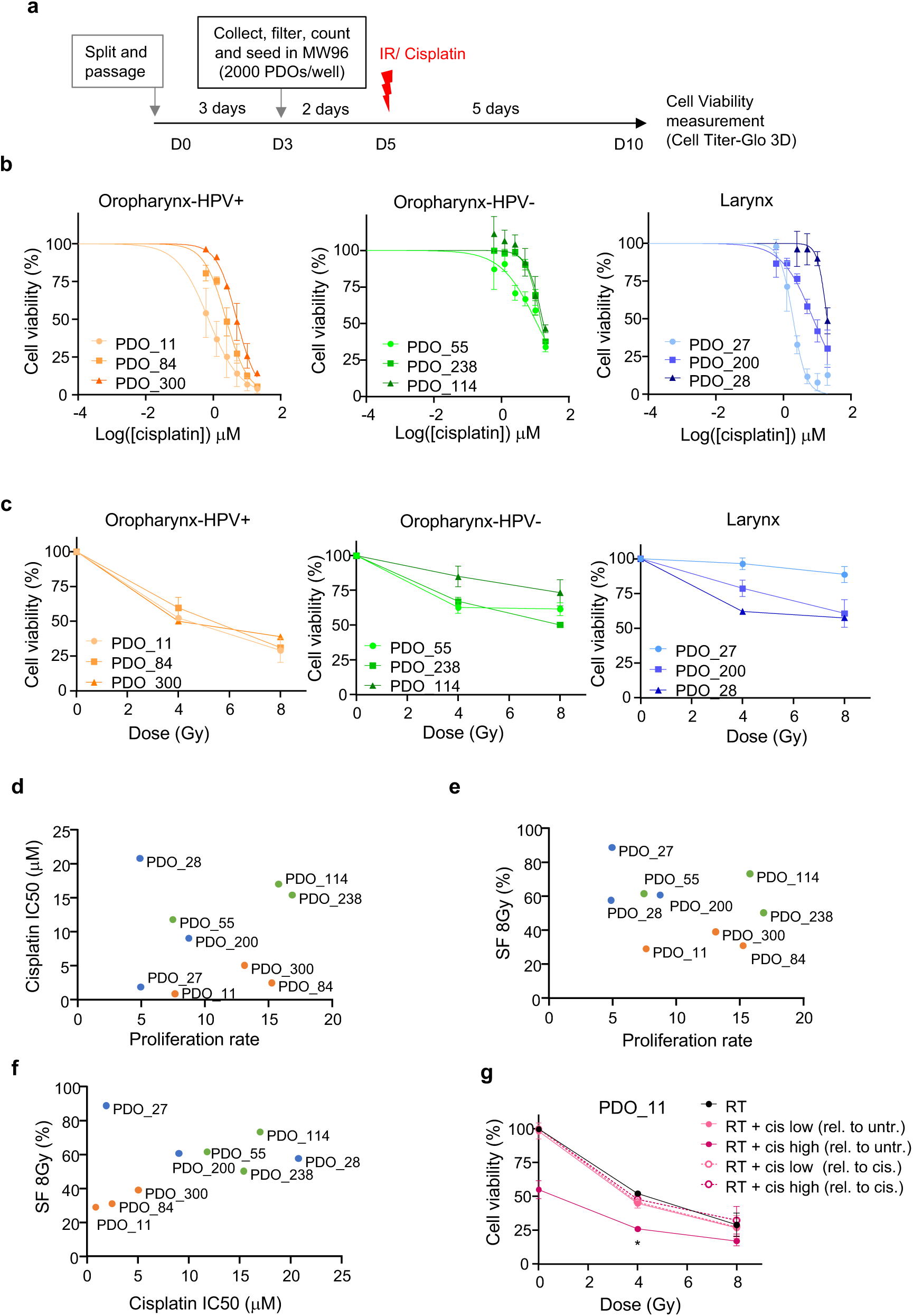
Differential response of HNSCC PDOs to cisplatin and radiation. **a)** Schematic overview of the treatment protocol. **b)** Cisplatin dose-response curves for the indicated organoid models derived from HPV-positive oropharyngeal (left), HPV-negative oropharyngeal (middle) and laryngeal tumors (right). Organoid cell viability is shown as percentage relative to untreated controls. Data represent the mean of two biological replicates; Error bars indicate standard error of the mean (SEM). **c)** Organoid viability (percentage relative to untreated controls) following exposure to 4 or 8 Gy of ionizing radiation in the indicated PDO cultures. Data represent the mean of two biological replicates; Error bars indicate SEM. **d)** Correlation between proliferation rate (x-axis) and cisplatin sensitivity based on IC_50_ (y-axis). Spearman correlation: r = 0.117, *p* = 0.78. **e)** Correlation between proliferation rate (x-axis) and radiation sensitivity based on survival fraction (SF) at 8 Gy (y-axis). Spearman correlation: r = -0.25, p = 0.52. **f)** Correlation between cisplatin IC_50_ values (x-axis) and survival fraction at 8 Gy (y-axis). Spearman correlation: r = 0.35, *p* = 0.36. **g)** Relative viability of PDO_11 after exposure to radiation (4 or 8 Gy) alone or in combination with cisplatin at low (1 µM) or high (5 µM) doses. Additive effects were calculated by normalization to untreated controls (*solid lines*), and synergistic effects by normalization to cisplatin-only treatment (*dashed lines*). Data represent the mean of two independent experiments. Error bars indicate SEM. Two-way Anova, *p* = 0.0147. SF, survival fraction; rel, relative; untr, untreated; cis, cisplatin.

A similar pattern was observed following IR exposure, with HPV-positive PDOs showing the lowest organoid survival fractions, consistent with reported clinical responses in patients **(Figure 4c)**. No significant correlation was detected between cisplatin IC₅₀ values or radiation survival fraction and PDO proliferation rate **(Figure 4d-e and Figure S3a)**. These findings indicate that differences in treatment response are not solely explained by proliferative capacity but are more likely driven by intrinsic tumor features. A weak positive association was observed between cisplatin and radiation responses

(Spearman r = 0.35, *p* = 0.36) **(Figure 4f)**, although this relationship did not reach statistical significance. Nevertheless, this trend is consistent with the clinical use of chemotherapy response in induction chemotherapy (ICT) programs to stratify patients as responders or non-responders for subsequent treatment with CRT or surgery, respectively [12].

Because concurrent CRT is a standard treatment for locally advanced or recurrent HNSCC, selected PDO lines were further treated with IR (4 Gy and 8 Gy) combined with sublethal cisplatin concentrations (1 and 5 µM) **(Figure 4g and Figure S3b-c)**. Potentiation of the radiation effect was observed only with the higher cisplatin dose (5 µM), predominantly in PDOs that were intrinsically sensitive to cisplatin, mainly HPV-positive models. Combined treatment with IR and 5 µM cisplatin significantly reduced viability in PDO_11 and showed a similar trend in other HPV-positive PDOs (PDO_84 and PDO_300), as well as in two laryngeal PDO lines (PDO_200 and PDO_28) **(Figure S3c)**. In these models, cisplatin demonstrated a radiosensitizing effect consistent with clinical observations. However, the combined treatment produced primarily additive rather than synergistic effects **(Figure 4g)**.

### Transcriptomic landscape of HNSCC organoids reveals distinct molecular subtypes

To further characterize these PDOs at the molecular level, transcriptomic profiling was performed by bulk RNA sequencing in 20 selected organoid lines **(Figure 5 and S4)** Principal component analysis (PCA) did not reveal clustering according to anatomical site **(Figure 5a)**. Therefore, unsupervised hierarchical clustering was conducted using the 500 most variable genes to determine whether alternative shared features could explain sample grouping. To define the optimal number of clusters, the sum of squared errors (SSE) was calculated for 1–20 clusters. The elbow method indicated an optimal range between five and nine clusters. Based on this analysis, eight clusters were selected, grouping samples with a dendrogram distance <0.9 **(Figure S4a)**. Notably, clustering was independent of the expansion medium used to establish organoid cultures.

**Figure 5.**
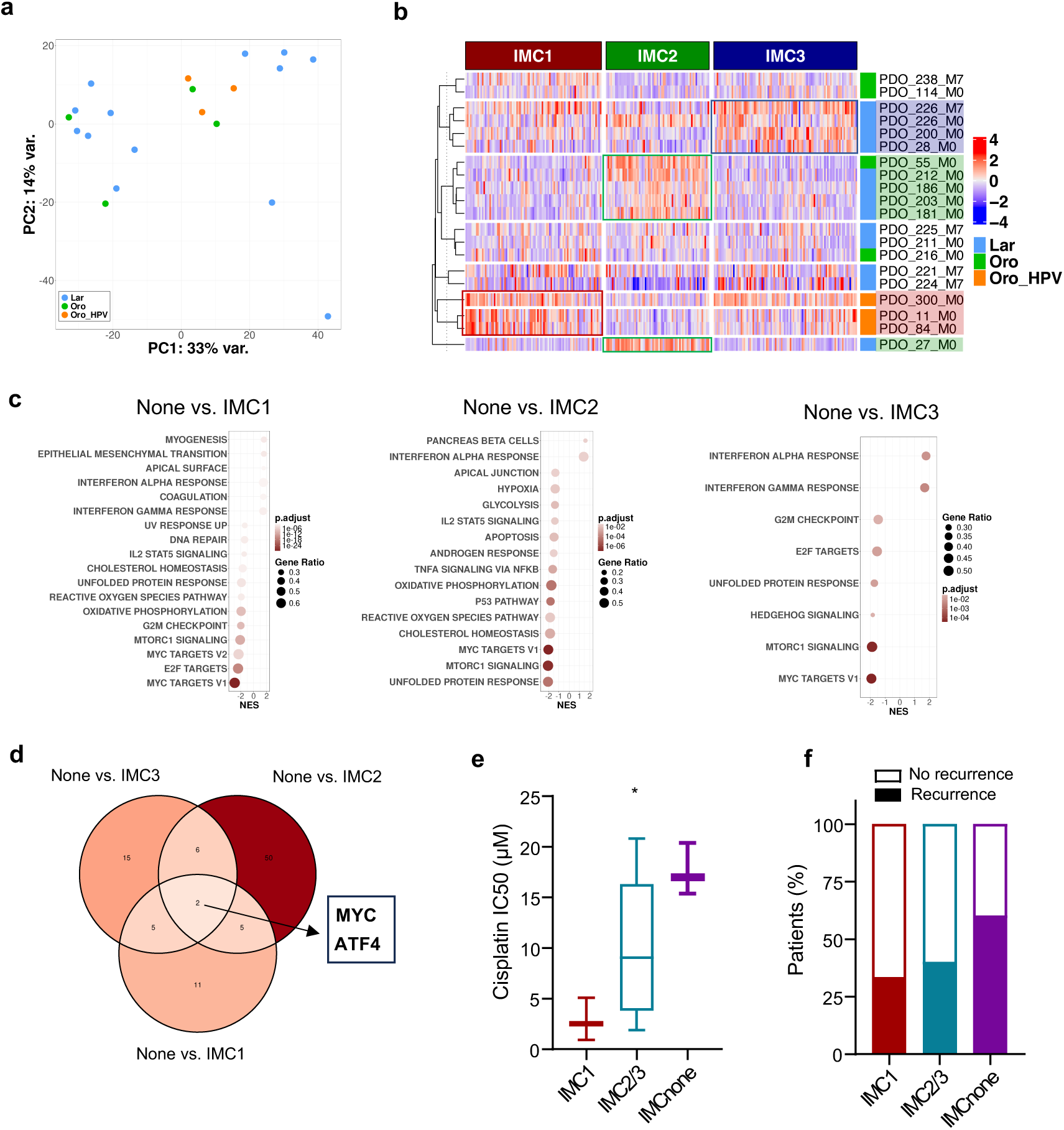
Molecular characterization of HNSCC PDOs by global transcriptomics. **a)** Principal component analysis (PCA) of bulk RNA-sequencing data from 20 PDOs, showing no segregation according to anatomical tumor site: larynx (Lar), oropharynx (Oro) and HPV-positive oropharynx (Oro_HPV). Samples are color-coded by tumor site (bottom left) and the percentage of variance explained by PC1 and PC2 is indicated. **b)** Heatmap displaying z-scored-normalized gene expression across PDO samples grouped into three molecular subtypes (IMC1, IMC2, and IMC3). Rows correspond to PDOs and columns to genes. Expression values are color-coded from blue (low expression) to red (high expression). PDOs are annotated by tumor site, as in **(a)**. Samples are ordered based on unsupervised hierarchical clustering performed using the 500 most variable genes (see **Figure S4a**). Selected gene clusters exhibiting subtype-specific expression patterns are highlighted (colored boxes), and subtype assignment is indicated by color-coded bars. **c)** Pathway enrichment analysis comparing PDOs not classified within IMC1-IMC3 (none) versus PDOs corresponding to IMC1 subtype (left), IMC2 (middle) or IMC3 subtype (right). Dot plots show normalized enrichment scores (NES), gene ratios, and adjusted *p* values. **d)** Venn diagram showing overlap of downregulated transcription factor target signatures in IMCnone PDOs compared with IMC1, IMC2, and IMC3. **e)** Cisplatin IC_50_ values across PDO molecular subtypes. Data are presented as box-and-whisker plots showing the meadian (*center line*), interquartile range (*box*) and whiskers extending to the minimum and maximum values. One-way Anova, * p*<* 0.05. **f)** Clinical correlation between PDO molecular classification and recurrence status of the primary tumors of origin.

We next assessed whether these clusters corresponded to previously described molecular HNSCC subtypes (IMC1, IMC2, and IMC3; [13]). Gene expression signatures defining these subtypes were visualized in a heatmap, with samples ordered according to the unsupervised clustering. As shown in both the heatmap **(Figure 5b)** and PCA **(Figure S4b)**, many organoids could be assigned to one of the reported subtypes. For example, HPV-positive PDOs (PDO_11, PDO_84, and PDO_300) were enriched for IMC1-associated genes, consistent with previous reports. Additional subsets of organoids were classified as IMC2 (PDO_55, PDO_181, PDO_186, PDO_203 and PDO_212) or IMC3 (PDO_28, PDO_200 and PDO_226), respectively **(Figure 5b)**.

Interestingly, a subset of organoids (PDO_211, PDO_216 or PDO_221 among others) did not clearly align with any of the established subtypes. To further characterize these samples, differential expression analysis was performed comparing them against organoids assigned to IMC1, IMC2, and IMC3. This analysis revealed a consistent downregulation of proliferation- and growth-related pathways, including mTORC1 signaling and MYC target genes **(Figures 5c and S4c–d)**, suggesting a more quiescent phenotype. Transcription factor target enrichment analysis further demonstrated reduced MYC and ATF4 activity across comparisons with all reported subtypes (Figures 5d and S4e).

Consistent with previous findings, IMC1 PDOs showed the highest sensitivity to cisplatin and were derived from patients with lower recurrence rates. In contrast, organoids corresponding to the putative new subtype(s) exhibited higher cisplatin IC_50_ values and were derived from tumors with higher recurrence rates, suggesting a more aggressive clinical behavior **(Figures 5e–f)**.

### In vivo tumorigenicity of HNSCC PDOs

Finally, organoid tumorigenicity was evaluated in vivo through xenotransplantation assays using immunodeficient mice. Organoids from two laryngeal PDO lines (PDO_200 and PDO_226) were subcutaneously injected into either nude mice or in a severely immunodeficient model (B-NDG), which lacks B, T and NK cells. PDO lines yielded macroscopically visible tumors although PDO_200 was the only one showing a clear exponential growth. Importantly, tumor growth was quite inefficient in nude mice, likely due to partial tumor clearance by the remaining immune cells present in these mice. In contrast, robust exponential tumor growth was observed in the ultra-immunodeficient model, predominantly for PDO_200, reaching nearly 800 mm^3^ in volume two months after transplantation **(Figure S5a)**. Histologically, H&E staining further revealed keratinization and stratification patterns that recapitulate various key features of the primary tumor of origin (Figure S5b).

## DISCUSSION

Here we report the establishment of the third largest collection of HNSCC PDOs and, to our knowledge, the largest series enriched in pharyngeal and laryngeal SCC (n > 50). This is particularly relevant as the largest previously reported HNSCC PDO collections were both predominantly composed of oral cavity tumors [9, 11]. Our overall success rate for long-term PDO expansion (35%) is consistent with those previous reports (30%-35%) [9, 11]. In agreement with previous work [11], we observed improved efficiency of organoid establishment using M7 medium compared with the original HN (M0) medium. Importantly, this improvement was not accompanied by major morphological or transcriptomic differences, indicating that enhanced culture conditions do not substantially alter the intrinsic molecular identity of the models.

Given the potential incorporation of PDOs into precision oncology programs, it is of major importance to determine whether specific clinical or tumor-related variables influence their successful establishment. In contrast to recent data from Dutch and German HNSCC PDO biobanks reporting higher establishment success in younger patients [9, 14], we did not observe any association between patient age and organoid formation efficiency. However, HPV positivity emerged as an independent positive predictor of PDO establishment. This observation has not been consistently reported in previous studies, likely due to differences in anatomical site composition. Thus, our analysis was restricted to oropharyngeal tumors (where HPV infection plays a relevant role), whereas earlier comparisons grouped multiple anatomical locations, predominantly oral cavity tumors [9, 14]. The higher establishment rate of HPV-positive tumors may reflect their lower tumor mutational burden, reduced chromosomal instability, and closer resemblance to non-transformed epithelial tissues. Across epithelial systems, organoid formation efficiency is frequently higher in healthy tissues than in tumor samples, and overgrowth of normal epithelial organoids has been described as a limitation for the expansion of tumor-derived organoids in certain tissues such a lung cancer [15] or prostate tumors [16]. Among all parameters analyzed, prior treatment with CT and/or RT emerged as the only independent negative predictor of PDO formation and expansion. While this finding contrasts with a previous report in HNSCC [14], it is consistent with other study also in HNSCC PDOs [9], and remained independent of the culture medium used, reinforcing its biological relevance. A plausible explanation for this negative impact is treatment-induced impairment of tumor cell fitness and stemness potential following genotoxic stress. A similar effect has been observed in primary bronchial epithelial cells, where organoid formation capacity is strongly reduced upon IR [17]. Analogously, radiotherapy-induced senescence has been shown to reduce organoid formation efficiency in salivary gland models [18]. Interestingly, this finding poses a challenge for the generation of *bona fide* therapy-resistant PDO models, particularly for studying acquired resistance mechanisms.

A distinctive feature of HNSCC PDOs is their derivation from stratified squamous epithelium. We observed a strong concordance between the morphological features and differentiation grade of PDOs and those of their tumor of origin. Although tumor grade is not currently use as a staging criterion in HNSCC, emerging evidence indicates that it may independently associate with immunotherapy response in recurrent and metastatic disease [19]. In the minority of cases in which PDOs did not fully recapitulate the differentiation status of the parental tumor, several factors may explain this discrepancy, including intratumoral heterogeneity, sampling bias during tissue acquisition, or the selective expansion of specific organoid-forming clones during *in vitro* culture.

Transcriptomic profiling further demonstrated that our PDO collection captures the molecular heterogeneity of HNSCC. Using a recently described IMC classifier [13], most PDOs were assigned to established molecular subtypes. However, a subset did not match any of the previously defined categories, suggesting the potential existence of at least one additional subtype. This putative new molecular subtype is characterized by reduced MYC and mTOR signaling, along with decreased expression of MYC target genes and diminished ATF4-related transcriptional programs. Given the central role of MYC and mTOR in anabolic metabolism, cell growth and proliferation, this transcriptional signature is consistent with a less proliferative phenotype [20-22]. Similarly, reduced ATF4 signaling, a key mediator of the unfolded protein response, suggests lower levels of proteotoxic stress [23]. At first glance, such a molecular profile might indicate a more indolent tumor biology and a potentially favorable prognosis. However, our functional data indicate that PDOs within this subtype exhibit increased resistance to cisplatin and are associated with higher recurrence rates. This apparent paradox may be explained by a more quiescent-like phenotype. Reduced MYC and mTOR signaling has been widely linked to cellular quiescence and dormancy [24-26], biological states that are typically less sensitive to cytotoxic agents targeting actively proliferating cells. Thus, rather than representing a biologically favorable subtype, these tumors may harbor intrinsic chemoresistance driven by reduced proliferative dependency.

Cisplatin-based chemoradiation remains a standard treatment for advanced HNSCC. In our functional assays, the combination of cisplatin and radiation enhanced tumor cell killing in a subset of PDOs, supporting the established clinical use of cisplatin as a radiosensitizer in the clinic [27]. However, our data suggest predominantly additive rather than synergistic effects, consistent with other independent organoid-based studies [11, 28]. A limitation shared across these studies, including ours, is the use of single-dose radiation rather than fractionated regimens that more closely mimic clinical protocols.

PDOs have been proposed as translational platforms to assess treatment responsiveness. Several observations from this study support their potential future use in functional precision medicine. HPV-positive PDOs were consistently more sensitive to cisplatin and/or radiation. This finding aligns with clinical evidence supporting HPV status as an independent favorable prognostic factor in oropharyngeal cancer [29], and is further supported by experimental models showing that HPV E6/E7 expression enhances radiosensitivity [8].

Additionally, we observed a positive association between cisplatin and radiation responses, supporting the clinical use of cisplatin response as a predictor of RT responsiveness in ICT programs for organoid preservation in laryngeal tumors [12]. Notably, the most cisplatin-resistant PDO in our cohort was derived from a recurrent tumor that relapsed six months after CRT, providing a clinically meaningful example of concordance between *ex vivo* drug response and patient outcome. Although preliminary, this observation highlights the potential utility of PDOs for functional assessment of treatment responsiveness.

## CONCLUSION

Overall, our data support the value of HNSCC PDOs as a robust 3D translational platform that faithfully captures key morphological, molecular, and functional features of the tumors from which they originate. Importantly, our data identify clinically meaningful determinants of organoid establishment, including the strong negative impact of prior chemo- and/or radiotherapy, and reveal a potential additional molecular subtype characterized by reduced MYC/mTOR signaling and increased cisplatin resistance.

Moreover, the organoids reproduced heterogeneous responses to standard treatments such as cisplatin and radiation, supporting their utility for modeling treatment response and resistance. Overall, these results highlight the potential of HNSCC PDOs as valuable platforms for studying tumor biology, identifying therapeutic vulnerabilities, and advancing functional precision oncology, although prospective studies using pre-treatment biopsies will be necessary to validate their predictive clinical value.

## LIST OF ABBREVIATIONS

CK13: Cytokeratin 13
CRT: Chemoradiotherapy
CT: Chemotherapy
FFPE: Formalin-fixed paraffin embedded
H&E: Hematoxylin and eosin
HNSCC: Head and neck squamous cell carcinoma
HPV: Human Papillomavirus
ICT: Induction chemotherapy
IHC: Immunohistochemistry
IR: Irradiation
OS: Overall survival
PDO: Patient-derived organoid
SCC: Squamous cell carcinoma
RT: Radiotherapy
SEM: Standard error of the mean

## Availability of data and materials

The RNAseq dataset supporting the conclusions of this article are available at the GEO database with the following accession number: GSE325165. All data generated and analyzed during this study are included in this published article and its supplementary information files and are available from the corresponding authors on reasonable request.

## Authors’ contributions

JP.R. and M.A-F. conceived and designed the study. M.A-F., M.A-G., E.P-A., and B.dL-D. established the organoid models and performed functional assays. H.C-M. contributed to organoid generation, treatment assays, and PDO RNA isolation. B.G. performed transcriptomic data analysis and visualization. I.R-G. and A.B. contributed to organoid culture and functional assays, respectively. M.O-R. and M.R-S. performed the in vivo mouse experiments. D.C-T. conducted histological and immunohistochemistry assays, while A.A. and I.F-V. performed the histological evaluation. S.B-P., F.L., and JP.R. contributed to patient selection, sample collection, and clinical data management. S.A-T., F.H-P., R.R., and JM.G-P. contributed to experimental design and data interpretation. JP.R. and M.A-F. performed statistical analysis. M.A-F. wrote the manuscript and supervised the study. M.A-G., E.P-A., B.dL-D., H.C-M., R.R., JM.G-P., and JP.R. reviewed and edited the manuscript.

## Funding

This study was supported by grants PID2020-117236RB-I00, PID2024-158165OB-I00 and CNS2023-144473, funded by MICIU/AEI/ 10.13039/501100011033 and, by the “European Union NextGenerationEU/PRTR”; a grant from the Scientific Foundation of the Spanish Association Against Cancer (LABAE235202ALVA) and a grant from Fundación Alimerka.

Additional funding was provided by the Instituto de Salud Carlos III (ISCIII) through the grants PI22/00167, PI24/00398 and PI25/00107; CIBERONC (CB16/12/00390), Fundación Bancaria Cajastur-IUOPA; Universidad de Oviedo, Instituto de Investigación Sanitaria del Principado de Asturias (ISPA) and grant PID2022-142020OB-I00 funded by MICIU/AEI/FEDER. Also funded by the Government of the Principality of Asturias through the Agency for Science, Business Competitiveness and Innovation of the Principality of Asturias and co-financed by the European Union, through the Grants “Subvenciones para Grupos de Investigación de Organismos del Principado de Asturias para el Ejercicio 2024” (IDE/2024/000778 and IDE/2024/000741). E.P.-A and M.O.-R. are both recipients of a PFIS predoctoral fellowship from ISCIII (FI20/00064 and FI23/00037, respectively). M.A.G. and B.G. are recipients of Severo Ochoa predoctoral fellowships from the Principado de Asturias (BP21-205 and BP20-046, respectively). F.H.-P and S.A.-T. are both recipients of a Miguel Servet fellowship from ISCIII (CP24/00064 and CP23/0010, respectively). D.C. was supported by the Platform ISCIII Biomodels and Biobanks through the project PT23/0077, funded by the Instituto de Salud Carlos III (ISCIII) and co-funded by the European Union.

## Acknowledgements

We thank the staff of the Animal facility at the University of Oviedo and the Radiophysics Unit at Hospital Universitario Central de Asturias (HUCA) for their excellent technical support. We want to particularly acknowledge the collaboration of the Principado de Asturias BioBank (PT20/00161 and PT23/00077), financed jointly by Servicio de Salud del Principado de Asturias, Instituto de Salud Carlos III, and Fundación Bancaria Cajastur and integrated in the Spanish National Biobanks and Biomodels Network.

## Notes

### Competing Interest Statement

The authors have declared no competing interest.

