## Supplemental Tables S2-S3 and Figures S1-S5 for "PATIENT-DERIVED ORGANOIDS CAPTURE HISTOLOGICAL, MOLECULAR AND THERAPEUTIC HETEROGENEITY IN PHARYNGEAL AND LARYNGEAL SQUAMOUS CELL CARCINOMAS"

**Table S2. Associations between organoid establishment in HN (M0) medium and clinical factors.**

| Feature | N samples (%) | Organoid formation | | Organoid expansion | |
|---|---|---|---|---|---|
| | | **Yes (derivation rate)** | ***P* value** | **Yes (freezing rate)** | ***P* value** |
| **- Sex** | | | | | |
| Male | 132 (94) | 90(68) | 1† | 41 (31) | 0.722† |
| Female | 9 (6) | 6 (67) | | 2 (22) | |
| **- Age** | | | | | |
| Below median (≤ 68.4) | | 45 (63) | 0.279† | 21 (30) | 0.856† |
| Above median (> 68.4) | | 51 (73) | | 22 (31) | |
| **- T classification** (n=112) | | | | | |
| T1-T2 | 52 (46) | 28 (54) | **0.033#** | 12 (23) | 0.187# |
| T3 | 34 (30) | 25 (73) | | 12 (29) | |
| T4 | 26 (23) | 21 (81) | | 11 (42) | |
| **- N classification** (n=102) | | | | | |
| N0 | 56 (55) | 36 (64) | 0.674† | 19 (34) | 0.832† |
| N1-3 | 46 (45) | 32 (70) | | 14 (30) | |
| **- Disease stage** (n=91) | | | | | |
| I-II | 32 (35) | 20 (62) | 0.126# | 10 (31) | 0.500# |
| III | 18 (20) | 12 (67) | | 6 (33) | |
| IV | 41 (45) | 34 (83) | | 18 (44) | |
| **- Pathological grade** (n=125) | | | | | |
| G1 | 43 (34) | 27 (63) | 0.566# | 13 (30) | 0.870# |
| G2 | 63 (50) | 45 (71) | | 21 (33) | |
| G3 | 19 (15) | 14 (74) | | 7 (37) | |
| **- Anatomic site** | | | | | |
| Oropharynx | 33 (23) | 24 (73) | 0.083# | 8 (24) | 0.527# |
| Hypopharynx | 16 (11) | 7 (43) | | 4 (25) | |
| Larynx | 92 (65) | 65 (71) | | 31 (34) | |
| **- HPV status** (n=33)‡ | | | | | |
| Negative | 24 (73) | 15 (62) | **0.039†** | 4 (17) | 0.170† |
| Positive (p16+) | 9 (27) | 9 (100) | | 4 (44) | |
| **- Treatment** | | | | | |
| Treatment naïve | 99 (70) | 74 (75) | **0.004†** | 30 (30) | **0.038†** |
| Surgery | 17 (12) | 12 (71) | | 9 (53) | |
| RT/CRT/CT | 25 (18) | 10 (40) | | 4 (16) | |
| **- Sample type** | | | | | |
| Surgical specimen | 120 (85) | 80 (67) | 0.456† | 38 (32) | 0.610† |
| Office biopsy | 21 (15) | 16 (76) | | 5 (24) | |
| **Total Cases** | **141** | **96 (68)** | | **43 (30)** | |

### Chi-square and † Fisher's exact tests. ‡ only oropharyngeal cases with HPV status.

**Table S3. Associations between organoid establishment in M7 medium and clinical factors.**

| Feature | N samples (%) | Organoid formation | | Organoid expansion | |
|---|---|---|---|---|---|
| | | **Yes (derivation rate)** | ***P* value** | **Yes (freezing rate)** | ***P* value** |
| **- Sex** | | | | | |
| Male | 37 (95) | 26 (70) | 1.000† | 14 (38) | 0.528† |
| Female | 2 (5) | 2 (100) | | 0 (0) | |
| **- Age** | | | | | |
| Below median (≤ 68.4) | | 13 (65) | 0.731† | 8 (40) | 0.514† |
| Above median (> 68.4) | | 15 (79) | | 6 (31) | |
| **- T classification** | | | | | |
| T1-T2 | 16 (41) | 13 (81) | 0.471# | 6 (38) | 0.262# |
| T3 | 15 (38) | 11 (73) | | 7 (47) | |
| T4 | 8 (21) | 4 (50) | | 1 (13) | |
| **- N classification** | | | | | |
| N0 | 21 (55) | 14 (67) | 0.497† | 7 (33) | 0.750† |
| N1-3 | 18 (45) | 14 (78) | | 7 (39) | |
| **- Disease stage** (n=36) | | | | | |
| I-II | 12(33) | 9 (75) | 0.832# | 5 (42) | 0.435# |
| III | 7 (19) | 5 (71) | | 4 (57) | |
| IV | 17 (47) | 11 (65) | | 5 (29) | |
| **- Pathological grade** (n=35) | | | | | |
| G1 | 6 (17) | 5 (83) | 0.540# | 2 (33) | 0.836# |
| G2 | 16 (46) | 10 (63) | | 6 (38) | |
| G3 | 13 (37) | 10 (77) | | 6 (46) | |
| **- Anatomic site** | | | | | |
| Oropharynx | 13 (33) | 10 (77) | 0.239# | 6 (46) | 0.548# |
| Hypopharynx | 5 (13) | 2 (40) | | 1 (20) | |
| Larynx | 21 (54) | 16 (76) | | 7 (33) | |
| **- HPV status** (n=13)‡ | | | | | |
| Negative | 6 (46) | 3 (50) | 0.070† | 2 (33) | 0.592† |
| Positive (p16+) | 7 (54) | 7(100) | | 4 (57) | |
| **- Treatment** | | | | | |
| Treatment naïve | 32 (82) | 26 (81) | **0.001**† | 14 (44) | 0.092† |
| Surgery | 2 (5) | 2 (100) | | 0 (0) | |
| RT/CRT/CT | 5(13) | 0 (0) | | 0 (0) | |
| **- Sample type** | | | | | |
| Surgical specimen | 30 (77) | 22 (73) | 0.693† | 11 (37) | 1.000† |
| Office biopsy | 9 (23) | 6 (67) | | 3 (33) | |
| **Total Cases** | **39** | **28 (72)** | | **14 (36)** | |

### Chi-square and † Fisher's exact tests. ‡ only oropharyngeal cases with HPV status.

#### Supplementary Figures

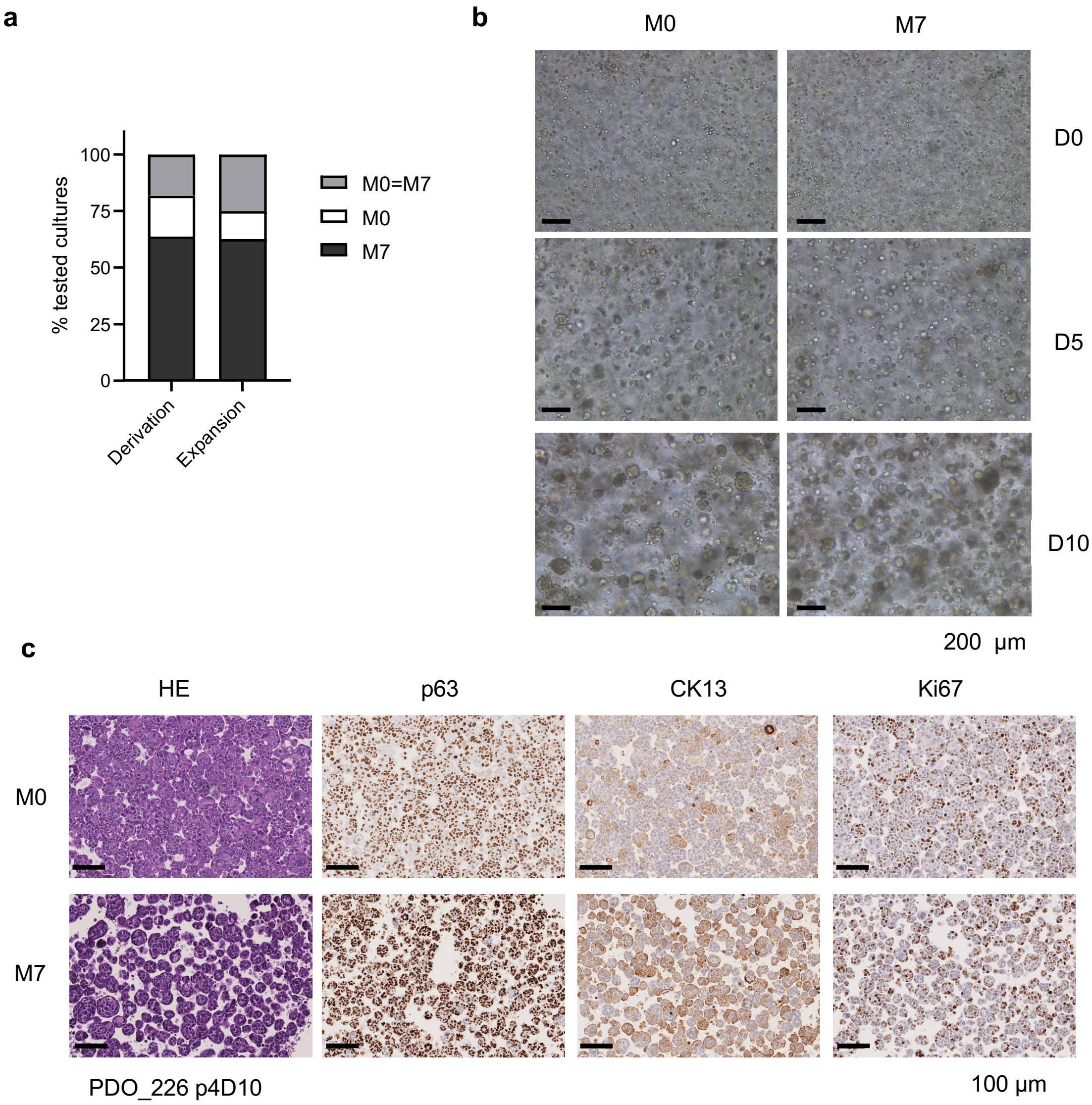


**Figure S1. Comparison of PDO establishment in HN (M0) versus M7 expansion media.**

**a)** Bar graph displaying the percentage of organoid models established and expanded with similar efficiency in both media (gray, M0 =M7), with improved growth in HN/M0 medium (white, M0 > M7), or with better performance in M7 medium (black, M7> M0). **b)** Representative bright-field images of PDO culture established and grown in each medium. **c)** Immunohistochemical analysis of a representative PDO model cultured under both conditions.

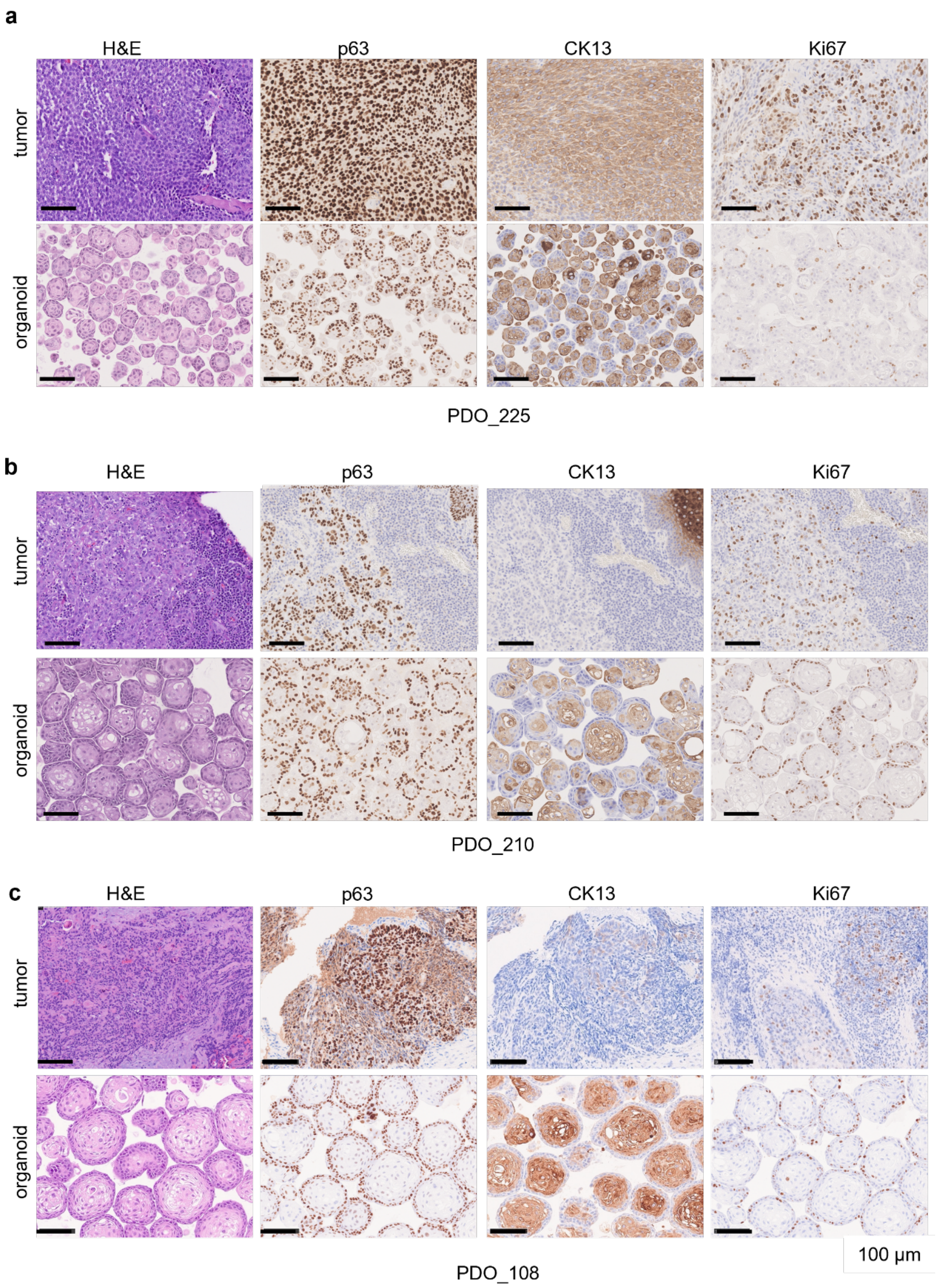


**Figure S2. Morphological evaluation of HNSCC tumors and matched PDOs showing partial histological concordance.**

Representative images of H&E staining and immunohistochemical detection of p63 (basal marker), CK13 (squamous differentiation marker) and Ki-67 ( proliferation marker) stainings in three HNSCC tumors and their matched PDOs, illustrating low or partial correlation between primary tumors and organoids: **a)** well-differentiated organoid versus poorly differentiated primary tumor (PDO_225, *top panels*); **b)** more differentiated and heterogeneous organoid versus moderately to poorly differentiated tumor areas (PDO_210, *middle panels*); and **c)** well-differentiated organoid versus heterogeneous, moderately to poorly differentiated tumor (PDO_108, *bottom panels*). Scale bars, 100 µm.

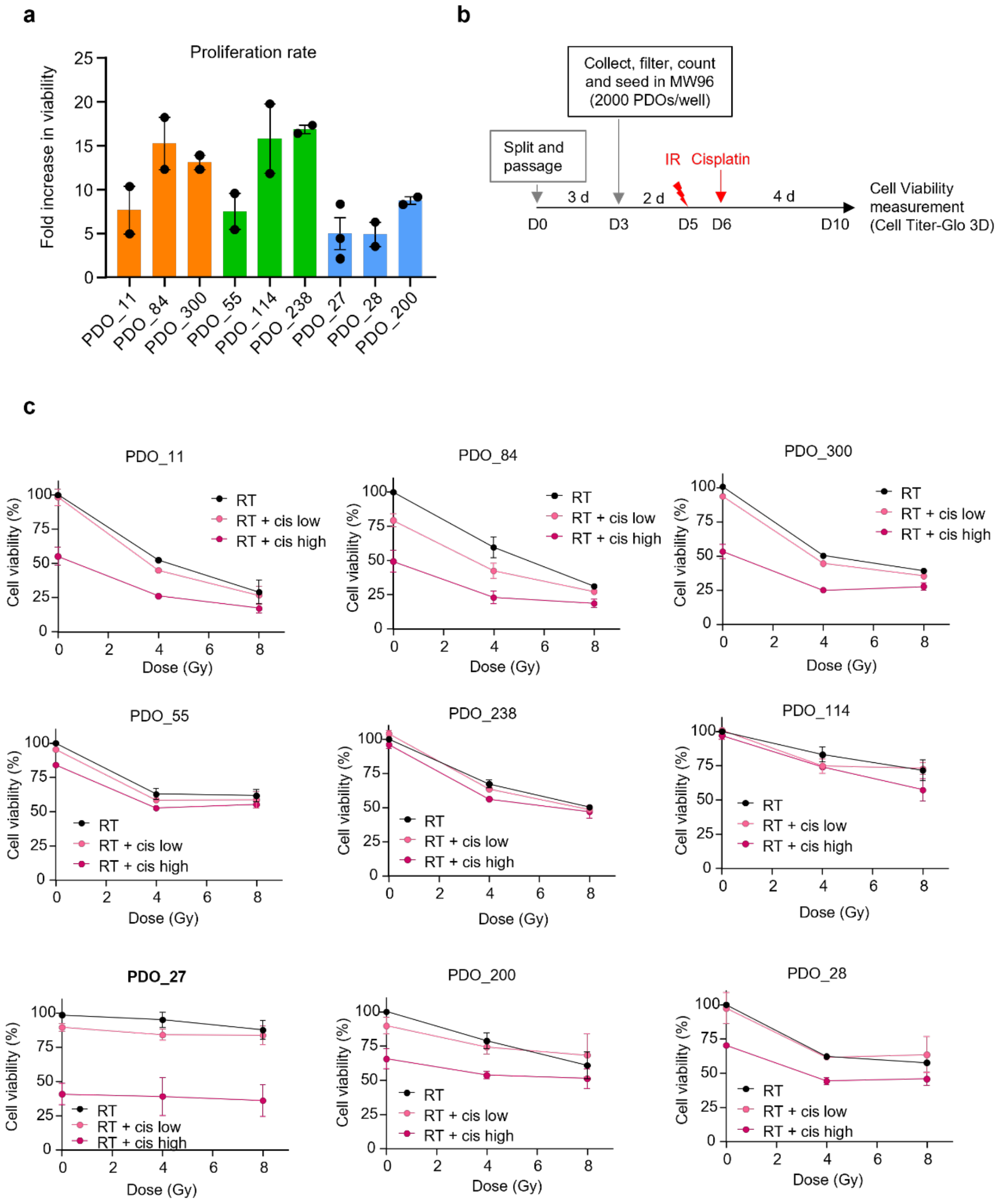

**Figure S3. Response of HNSCC PDOs to chemoradiotherapy.**

**a)** Proliferation rate of the indicated PDOs determined from the increase in organoid viability over 7 days in the absence of treatment. Data represent the mean of two independent experiments. Error bars indicate SEM. **b)** Schematic overview of the treatment protocol for chemoradiotherapy. **c)** Relative viability of the indicated PDOs after exposure to radiation (4 and 8 Gy) alone or in combination with cisplatin at low (1 μM) or high (5 μM) doses. Potential additive effects were calculated by normalization to untreated controls. Data represent the mean of two independent experiments. Error bars indicate SEM. Two- way Anova, $p > 0.05$ (not significant). cis, cisplatin.

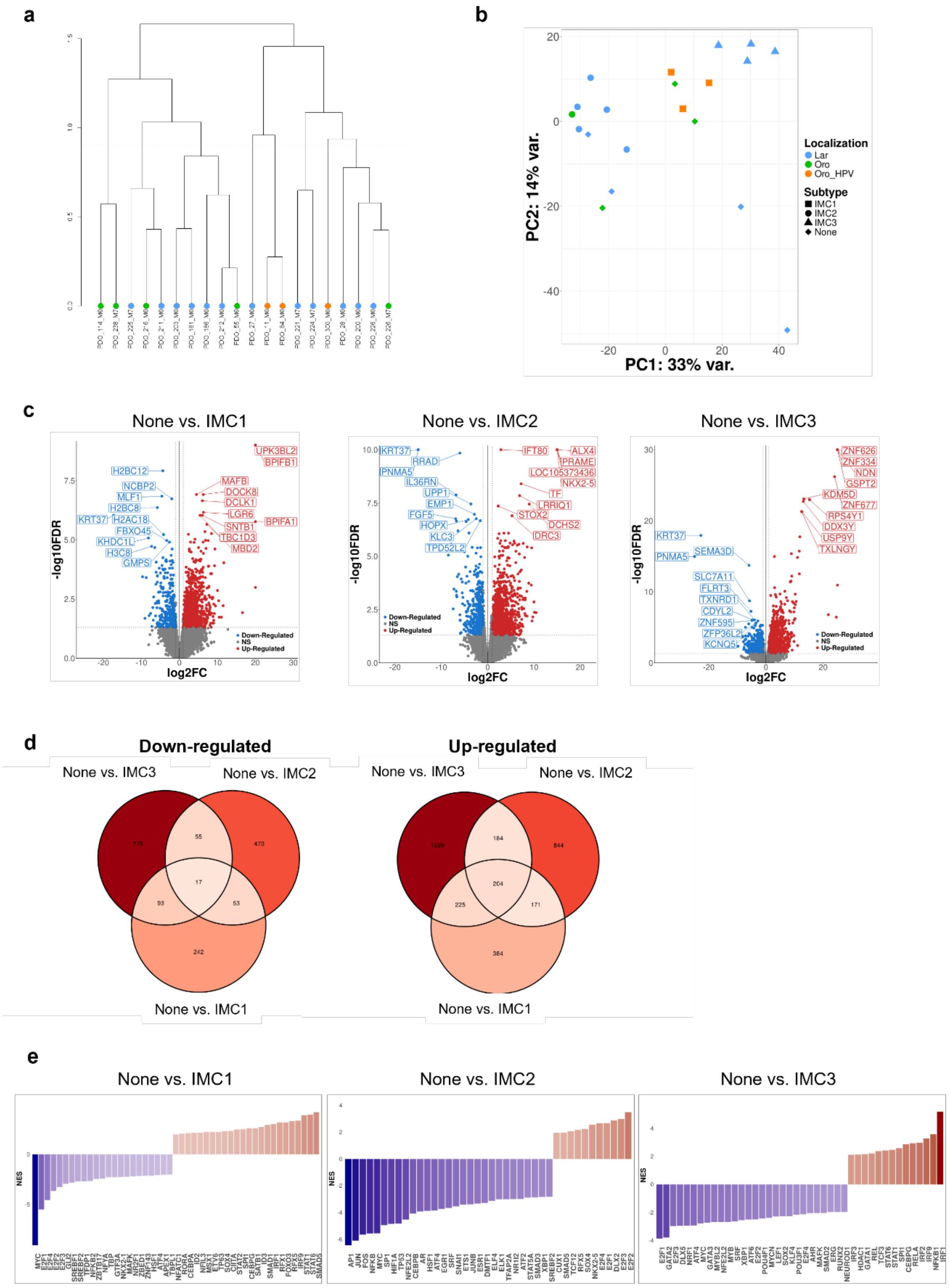


**Figure S4. Transcriptomic analysis of HNSCC PDOs.**

**a)** Unsupervised hierarchical clustering of 20 PDOs using the 500 most variable genes. The dendrogram shows sample relationships based on transcriptomic similarity. **b)** Principal component analysis (PCA) of bulk RNA-sequencing data. Samples are colored according to anatomical location and shaped according to molecular subtype (IMC1,

IMC2, IMC3, or IMCnone). Variance explained by PC1 and PC2 is indicated. **c)** Volcano plots showing differential gene expression analyses comparing IMCnone PDOs versus IMC1, IMC2, and IMC3 subtypes. Red dots represent significantly upregulated genes,and blue dots represent significantly downregulated genes. Selected top differentially expressed genes are annotated. **d)** Venn diagrams depicting overlap of significantly downregulated (*left*) and upregulated (*right*) genes in IMCnone PDOs relative to IMC1, IMC2, and IMC3 subtypes. **e)** Distribution plots of transcription factor target signatures differentially expressed in IMCnone versus each IMC subtype Bars represent normalized enrichment scores (NES) for individual transcription factors ordered by fold change. Red bars represent significantly upregulated transcription factor signatures, whereas blue bars indicate significantly downregulated signatures.

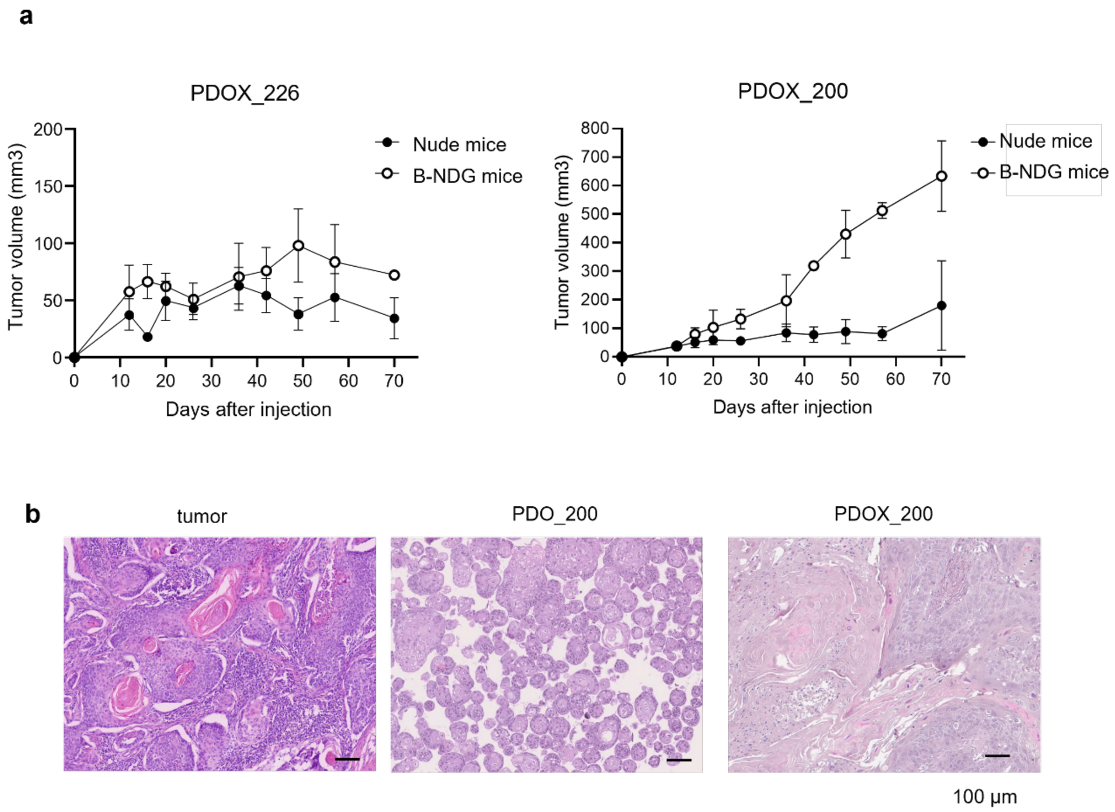


**Figure S5. Characterization of HNSCC PDO xenograft (PDOX).**

**a)** Tumor growth curves (mean +/- SEM) of PDOX tumors in nude or B-NDG mice after transplantation of PDO_226 (*left panel*) or PDO_200 (*right panel*) organoids. **b)** Representative H&E staining of the corresponding tumor of origin (*left panel*), derived PDO (*middle panel*) and PDOX grown in B-NDG mice (*right panel*). Scale bars, 100 µm.
